# Viral hijacking of the nucleolar DNA-damage response machinery: a novel mechanism to regulate host cell biology

**DOI:** 10.1101/219071

**Authors:** Stephen M. Rawlinson, Tianyue Zhao, Ashley M. Rozario, Christina L. Rootes, Paul J. McMillan, Anthony W. Purcell, Amanda Woon, Glenn A. Marsh, Kim G. Lieu, Lin-Fa Wang, Hans J. Netter, Toby D. Bell, Cameron R. Stewart, Gregory W. Moseley

## Abstract

Recent landmark studies indicate that nucleoli play critical roles in the DNA-damage response (DDR) *via* interaction of DDR machinery including NBS1 with nucleolar Treacle protein, a key mediator of ribosomal RNA (rRNA) transcription and processing, implicated in Treacher-Collins syndrome. Here, using proteomics, confocal/super-resolution imaging, and infection under BSL-4 containment, we present the first report that this nucleolar DDR pathway is targeted by infectious pathogens. We find that Treacle has antiviral activity, but that matrix protein of Henipaviruses and P3 protein of rabies virus, highly pathogenic viruses of the order *Mononegavirales*, interact with Treacle and inhibit its function, thereby silencing rRNA biogenesis, consistent with mimicking NBS1-Treacle interaction during a DDR. These data identify a novel mechanism for viral modulation of host cells by appropriating the nucleolar DDR; this appears to have developed independently in different viruses, and represents, to our knowledge, the first direct intra-nucleolar function for proteins of any mononegavirus.

---

The DNA damage response (DDR) comprises a complex network of pathways that monitor and repair damage to genomic DNA to prevent deleterious mutations^1^. The mechanisms underlying the DDR are only partially resolved, but recent studies implicated nucleoli as having critical roles^2-5^. The canonical function of nucleoli is ribosome biogenesis, where they support ribosomal RNA (rRNA) synthesis, processing, and assembly into pre-ribosomal subunits; however, numerous studies have indicated that nucleoli are highly multifunctional and dynamic structures involved in processes including stress responses, cell-cycle regulation and signal recognition particle assembly^6^. These functions derive from a large nucleolar proteome, by which nucleoli are considered to act as integrators of complex cellular signals, the full extent and mechanisms of which are only beginning to be understood^6^.

Recent studies of the roles of nucleoli in stress responses have identified Treacle protein, a nucleolar regulator of rDNA transcription and pre-rRNA processing that localizes to subnucleolar compartments (referred to here as Treacle-enriched puncta), as a critical mediator of global rRNA silencing that is induced by the DDR^2,4,7,8^. Specifically, DDR-induced rRNA silencing requires Nijmegen Breakage Syndrome 1 (NBS1) protein, a key effector of the DDR that forms part of the MRN complex^1,2,4^. NBS1 interacts with Treacle and, during a DDR, accumulates in Treacle-enriched puncta to induce rRNA silencing^2,4^. Under normal conditions, Treacle maintains basal levels of rRNA biogenesis such that depletion of Treacle expression results in reduced rRNA synthesis^4,7^. During the DDR, the extent of rRNA inhibition observed is equivalent to that following Treacle depletion, and induction of the DDR in cells depleted for Treacle causes no additional inhibition^4^. Thus, DDR-induced silencing of rRNA synthesis is Treacle-dependent, and the data appear consistent with a model whereby Treacle’s normal function in rRNA biogenesis is inhibited *via* the NBS1-Treacle complex; however the precise mechanism has not been confirmed^2,4^. Notably, reduced Treacle expression and consequent rRNA biogenesis is associated with the genetic disorder Treacher-Collins Syndrome (TCS), a severe craniofacial developmental disorder wherein mutations of the Treacle-encoding TCOF1 gene account for the majority of cases^9^. Haplo-insufficiency of Treacle is thought to result in insufficient ribosome biogenesis in highly proliferative neuroepithelial cells during development, leading to nucleolar stress and activation of apoptosis^10^.

Other than its roles in genetic disorders such as TCS, neurodegenerative diseases^11^, and cancers^12^, the nucleolus is targeted by proteins expressed by diverse viruses, potentially enabling viral modulation of intra-nucleolar processes controlling host cell biology^13,14^. However, this aspect of viral biology remains poorly characterized, particularly with respect to viruses of the order *Mononegavirales*, which comprises non-segmented negative-strand RNA viruses including the highly pathogenic and medically significant Hendra (HeV) and Nipah (NiV) viruses (genus *Henipavirus*, family *Paramyxoviridae*), rabies virus (RABV; genus *Lyssavirus,* family *Rhabdoviridae*), Ebola virus, measles virus, and mumps virus. Almost all mononegaviruses mediate transcription, replication and assembly exclusively within the cytoplasm, but several recent reports indicate that certain proteins of mononegavruses can localize to the nucleolus and bind to specific nucleolar proteins^15-17^.

The henipavirus matrix (M) protein is perhaps the best characterized of the nucleolar proteins expressed by mononegaviruses. The key henipavirus members are HeV and NiV, which are zoonotic viruses that have emerged from bat reservoirs to cause multiple outbreaks in humans and domesticated animals, with mortality rates between 40% - 100%, and no human vaccine or therapeutic available^18^. Henipavirus M protein plays essential roles in virus particle assembly in the cytoplasm and budding at the plasma membrane, which are broadly conserved among paramyxoviruses and other mononegaviruses^19,20^. However, M protein also enters the nucleus and accumulates within nucleoli in NiV- and HeV-infected cells^17,21^. Recent studies reported a requirement for M protein to traffic to the nucleolus prior to fulfilling its role in budding^17,22^. Furthermore, certain proteomic data suggest that M protein can interact with multiple nucleolar factors, supporting the possibility of an intranucleolar role^22,23^. However, it has also been proposed that the principal role of nuclear/nucleolar localization of proteins of mononegaviruses might be to sequester inhibitory proteins from the cytoplasm during replication prior to budding,^13,20,24^ and, importantly, no direct nucleolar function has been reported for M protein or for nucleolar proteins expressed by any other mononegavirus; thus, the potential significance of such interactions remains unresolved.

Here, we show that M protein of henipaviruses as well as the distinct P3 protein of another mononegavirus, rabies virus, specifically target Treacle protein and inhibit rRNA biogenesis. The data indicate that M protein enters Treacle-enriched puncta and exploits the DDR-Treacle pathway by a mechanism consistent with mimicking DDR-activated NBS1. This identifies a novel viral strategy to subvert host cell biology that appears to be utilized by diverse viruses.

## RESULTS

### Hendra virus M protein accumulates in a subnucleolar compartment

Nucleoli comprise multiple compartments with discrete functions such that protein localization to particular subnucleolar regions enables different regulatory roles^25,26^. Although HeV M-protein is known to target the nucleolus^15,17,22^ its specific subnucleolar localization has not been described. To examine this, we used confocal laser scanning microscopy (CLSM) to image nucleoli in cells infected by HeV and immunostained for M protein (Fig. 1A), or in living cells transfected to express HeV M protein fused to green fluorescent protein (GFP-HeV M) with or without the nucleolar marker nucleolin (NCL), fused to mCherry (mCherry-NCL) (Fig. 1B). Rather than being diffuse within nucleoli, M protein accumulated strongly within discrete subnucleolar puncta in both infected and transfected cells, indicating an intrinsic capacity of M protein to interact with specific subnucleolar compartments, which is independent of other viral proteins.

**Figure 1.**
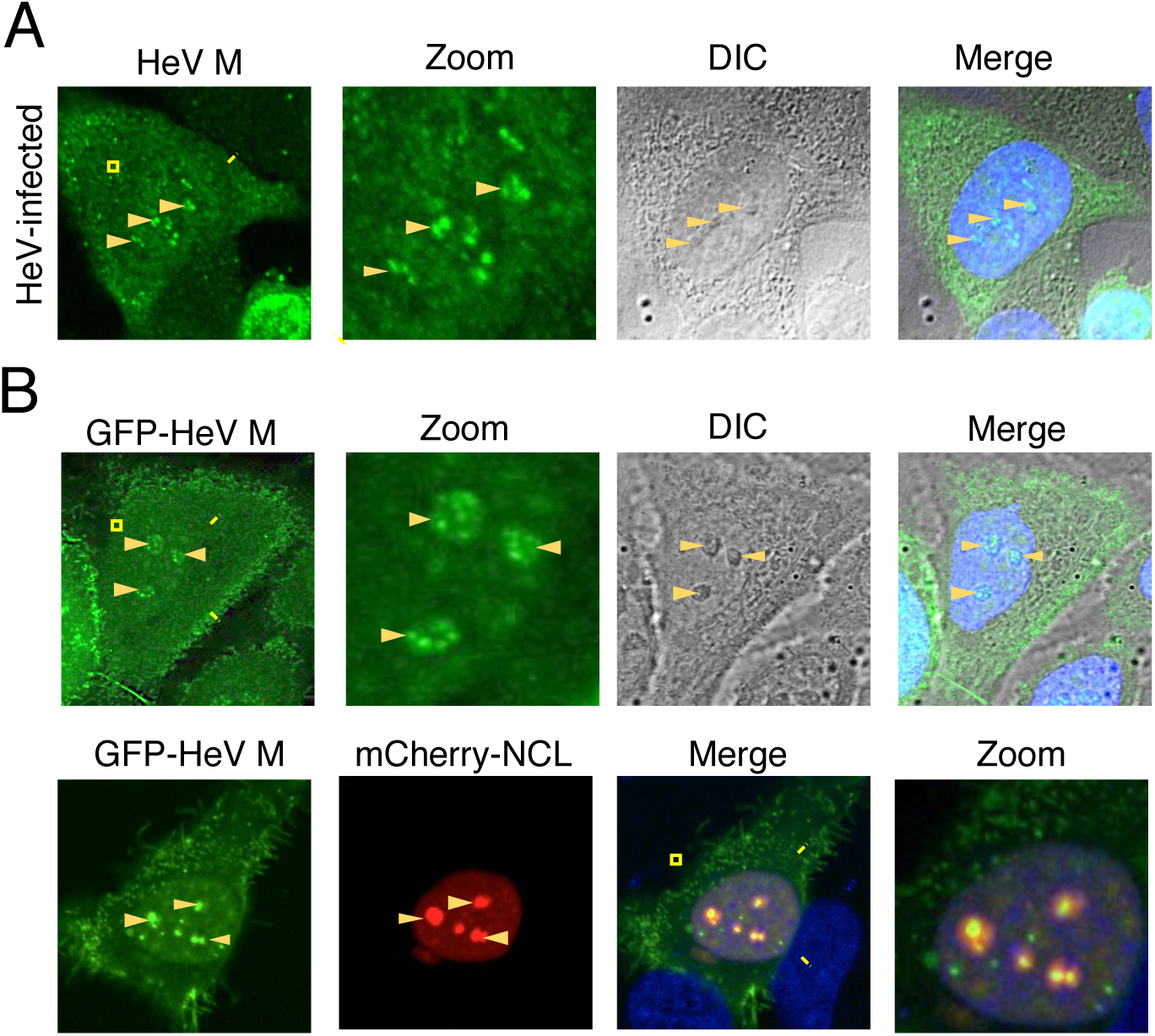
HeV M protein localizes to discrete subnucleolar compartments in HeV-infected cells and cells transfected to express HeV M protein. (A) HeLa cells infected with HeV (multiplicity of infection (MOI) of 5) were fixed 24 h post-infection (p.i.) and immunostained for HeV M protein before imaging by CLSM. (B) HeLa cells transfected to express GFP-HeV M protein alone (upper panel), or with mCherry-NCL (lower panel) were imaged live at 16 h post transfection (p.t.) by CLSM. Images are representative of ≥ 5 independent experiments, in each of which ≥ 10 fields of view were captured per condition. Arrowheads indicate nucleoli. Nuclei (DNA) were labeled using Hoechst 33342 (blue in merged images). Yellow boxes highlight regions of images magnified in the Zoom panel.

Fibrillarin (FBL) is a nucleolar protein that was identified by a genome-wide functional genomics screen to be required for HeV production, and reported to interact with M protein, although no function has yet been identified for this interaction^15^. We thus examined the subnucleolar localization of GFP-HeV M and red fluorescent protein-fused FBL (RFP-FBL) in living cells using super-resolution imaging (Fig. 2A). This confirmed localization of GFP-HeV M to subnucleolar puncta, but FBL was clearly excluded from these structures (Fig. 2A; Supplementary Movie 1). Fluorescence intensity profiles confirmed that the punctate regions enriched for GFP-HeV M were largely devoid of RFP-FBL (Fig. 2B). Thus, the localization of HeV M protein to subnucleolar puncta does not appear to involve interaction with FBL.

**Figure 2.**
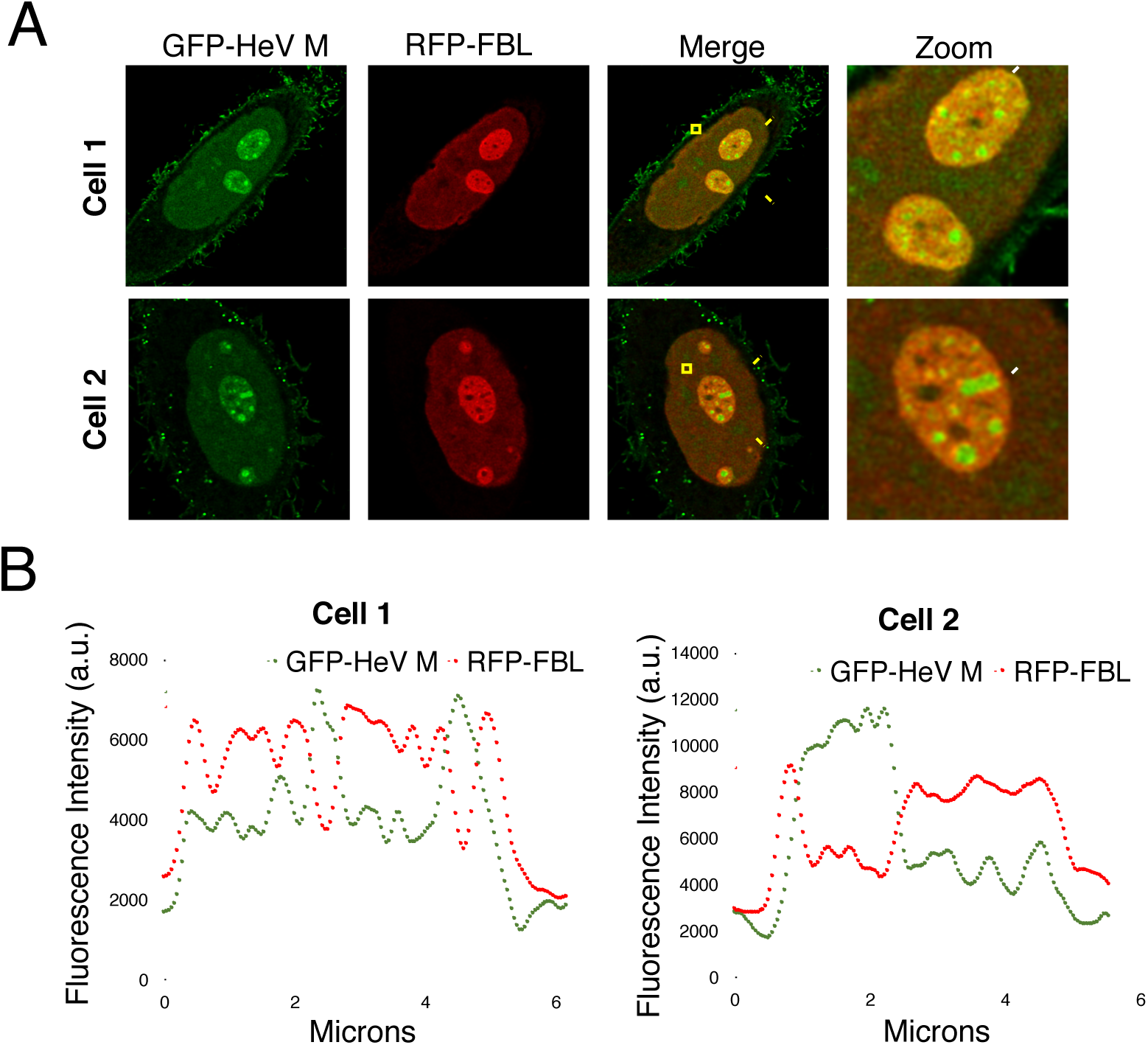
FBL is excluded from HeV M-containing sub-nucleolar puncta. (A) HeLa cells transfected to express GFP-HeV M and RFP-FBL were imaged live at 16 h p.t. using a Leica SP8 confocal microscope with Hyvolution to achieve super-resolution. Images are representative of cells in ≥ 5 fields of view. Yellow boxes highlight regions magnified in the Zoom panel. (B) Fluorescence intensity profiles generated from images in A; dotted lines in the Zoom panels indicate regions selected for profiling.

### Mutation of HeV M protein residue K258 to alanine abolishes subnucleolar puncta localization without impairing localization into nucleoli

Henipavirus M protein contains two nuclear localization sequences (NLSs) and two nuclear export sequences (NESs)^17,27^, but the requirements for nucleolar targeting are unknown. Since NLSs and nucleolar localization sequences (NoLSs) are often proximal or overlap^28^ we examined the impact on nucleolar localization of mutation of K258 to alanine (K258A), which was reported to inhibit nuclear localization of NiV M protein suggesting that K258 forms part of a C-terminal NLS^17,22^.

GFP-fused M proteins of both HeV and NiV formed clear subnucleolar puncta in ≥ c. 90% and 80%, respectively, of nucleoli examined (Fig. 3A, B). Mutation of K258 to A did not prevent accumulation of HeV or NiV M protein within nucleoli, but strongly impaired accumulation within subnucleolar puncta, such that no punctate localization was observed in nucleoli of cells expressing mutated HeV or NiV M protein (Fig. 3A, B). This suggested that mutation of K258 inhibits localization of M protein to puncta and, consistent with this, we were able to detect clear exclusion of K258A-mutated M protein from subnucleolar punctate structures (Fig. S1). Interestingly, although K258A mutation of NiV M protein resulted in a more cytoplasmic localization, as previously reported^17^, HeV M protein mutated at K258A remained largely nuclear (Fig. 3A). Thus, it appears that the requirements for subnucleolar localization are conserved between HeV and NiV M proteins, while those for nucleo-cytoplasmic localization differ, in spite of high sequence conservation between the proteins (c. 90% amino acid identity). Importantly, the finding that K258A mutation did not prevent nucleolar localization of HeV M protein, but resulted in a failure to accumulate within subnucleolar puncta, provided the opportunity to define the nature of the puncta and to examine the specific roles of punctate localization.

**Figure 3.**
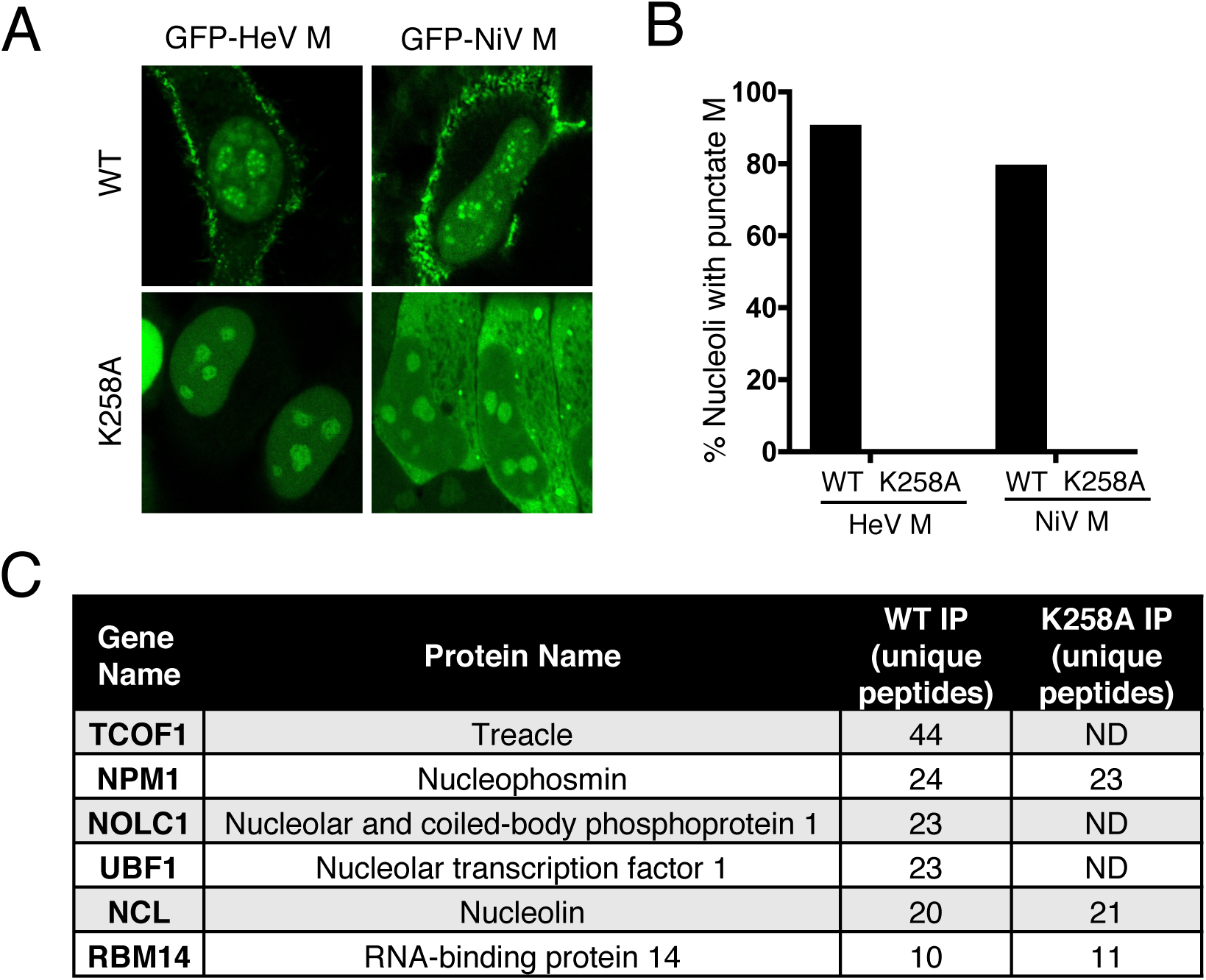
Mutation of residue K258 to alanine in HeV M protein inhibits localization to subnucleolar puncta and binding to a subset of nucleolar proteins. (A) HeLa cells transfected to express the indicated proteins were analyzed by live-cell CLSM. (B) Images such as those in (A) were used to determine the percentage of nucleoli with clear localization of M protein to subnucleolar puncta (≥ 447 nucleoli analyzed for each of wt and K258A-mutated M protein from 4 separate assays; ≥ 99 nucleoli analyzed for each of wt and K258A-mutated NiV M protein, 2 separate assays). (C) Nucleolar interactors of GFP-HeV M and GFP-HeV M-K258A identified by IP/MS; proteins are ranked by number of unique peptides identified. ND = not detected.

### HeV M protein associates with Treacle within Treacle-enriched puncta

To identify cellular factors associated with the subnucleolar localization of M protein we performed comparative immunoprecipitation/mass spectrometry (IP/MS) to compare the nucleolar protein interactome of wt and K258A-mutated GFP-HeV M. The most pronounced interactors of wt M protein, according to the number of unique peptides, included nucleolar proteins Treacle, nucleolar and coiled-body phosphoprotein 1 (NOLC1) and nucleolar transcription factor 1 (UBF1) (Fig. 3C). None of these proteins were detected in IP/MS of GFP-only controls (not shown) and, notably, were not detected in the interactome of K258A-mutated HeV M protein. Since wt and K258A-mutated HeV M protein interacted similarly with other nucleolar proteins, including nucleophosmin (NPM1), NCL and RNA-binding protein 14 (RMB14), it appeared that K258A specifically impacts interactions with a subset of nucleolar proteins, indicating that these interactors are relevant to punctate localization.

The most pronounced nucleolar interactor was Treacle, which is known to localize to subnucleolar puncta^29^, reminiscent of HeV M nucleolar localization (Figs. 1, 2), and so was selected for further analysis. To assess the impact of Treacle on the production of infectious HeV, we transfected cells with negative control (scr) siRNA or siRNA targeting Treacle, before infection with HeV. The specific depletion of Treacle expression (Fig. S2) resulted in a significant (p < 0.003) increase in virus production (Fig. 4A), indicating that Treacle has antiviral properties.

**Figure 4.**
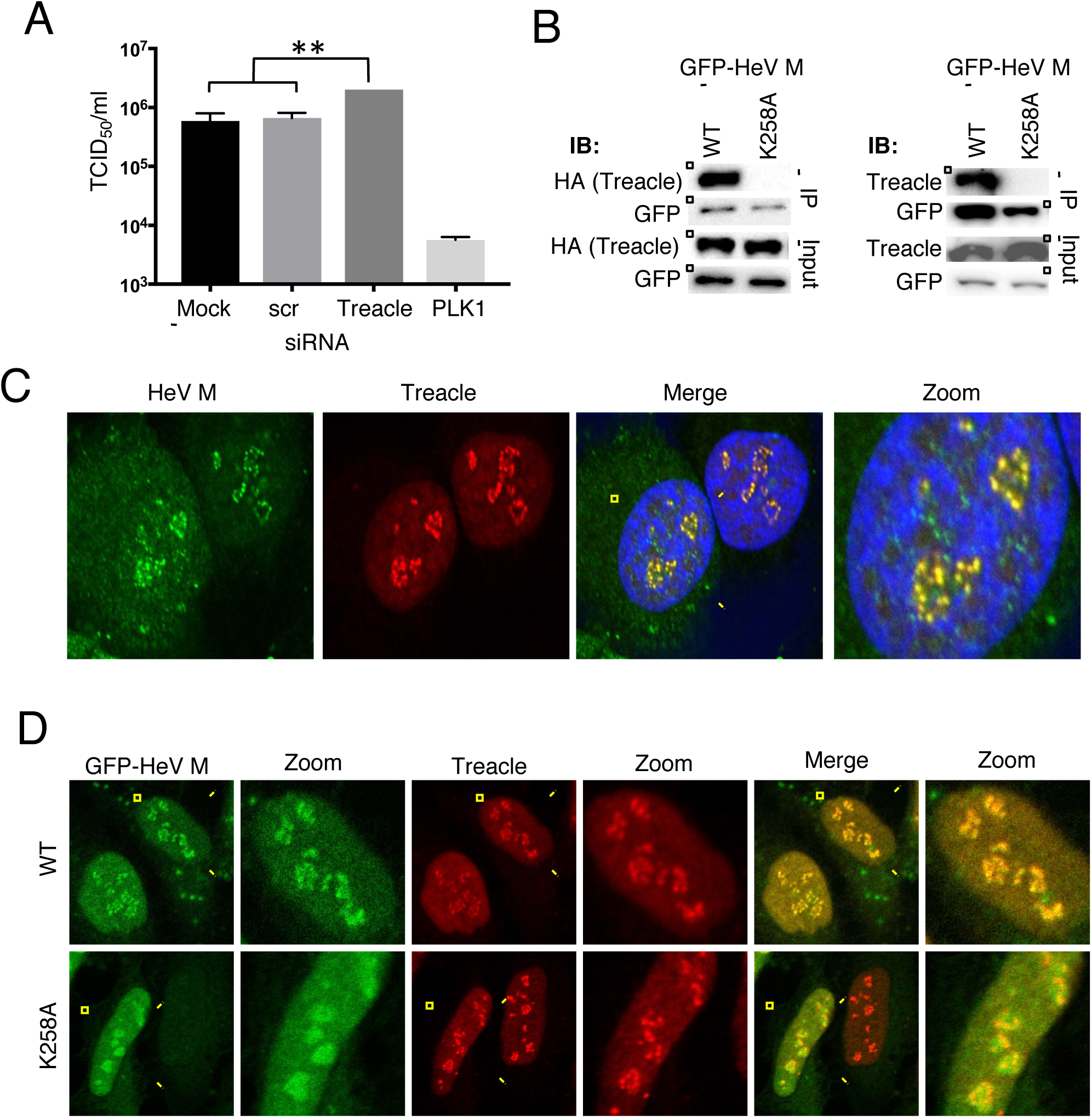
Treacle impacts HeV production and interacts with wt but not K258A HeV M protein. (A) HeLa cells were transfected with scrambled siRNA (scr), Treacle siRNA, or mock-transfected, before infection with HeV (MOI 0.5); siRNA for polo-like kinase 1 (PLK1) is a positive control known to inhibit HeV production^15^. HeV titer was measured at 48 h p.i. (mean TCID_50_/mL, ± SEM, n = 3). Statistical analysis used Student’s t-test; **, p < 0.001. (B) IPs of the indicated proteins from HEK-293T cells were analyzed by immunoblotting (IB) with the indicated antibodies; results are representative of 3 independent experiments. (C) HeV-infected cells were fixed 24 h p.i. and immunostained for HeV M protein and Treacle. Nuclei were labelled using Hoechst 33342 (blue in merged image). Images are representative of ≥ 20 fields of view, capturing ≥ 100 cells over 2 experiments. (D) HeLa cells transfected to express the indicated proteins were fixed and immunostained for Treacle; images are representative of > 30 fields of view over 3 experiments.

Using IP and immunoblot (IB) analysis, we further confirmed that GFP-HeV M interacts with endogenous Treacle and with co-transfected HA-Treacle, and that mutation of K258 prevents these interactions (Fig. 4B). Co-localization of HeV M with endogenous Treacle within the subnucleolar puncta was also confirmed by CLSM analysis of infected cells (Fig. 4C) and cells expressing GFP-HeV M (Fig. 4D upper panels). HeV M K258A did not accumulate with Treacle in puncta (Fig. 4D lower panels). Notably, siRNA knockdown of Treacle inhibited the accumulation of GFP-HeV M into puncta (Fig. 5A, B), with c. 95% of nucleoli in Treacle-depleted cells showing a diffuse nucleolar localization, and exclusion from puncta often detected, resembling the localization observed in cells expressing GFP-HeV M-K258A (Fig. 3A). Thus, it appears that M protein localizes to the previously described Treacle-enriched puncta^29^, within which it interacts with Treacle protein.

**Figure 5.**
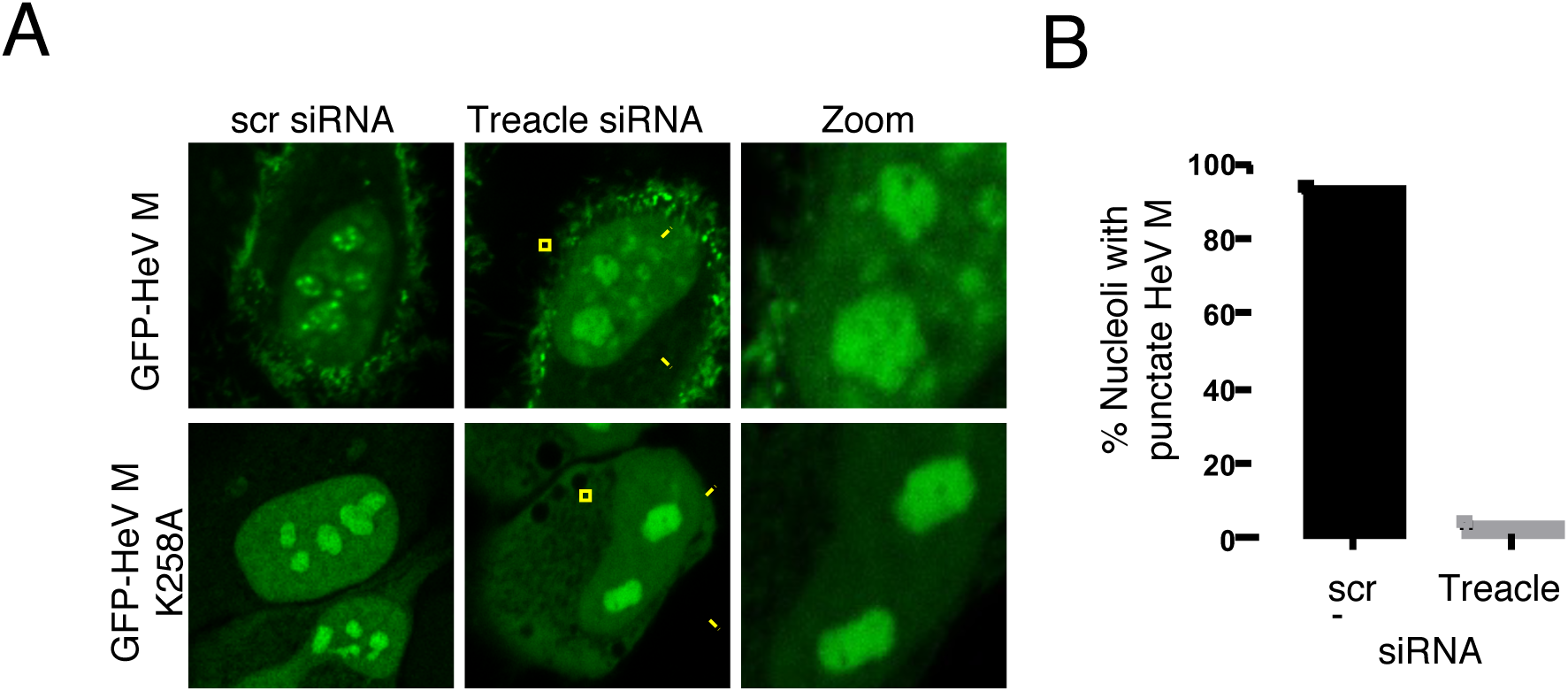
Punctate localization of HeV M is dependent on Treacle. (A) HeLa cells were transfected with scr siRNA or Treacle siRNA 3 days prior to transfection to express the indicated GFP-HeV M proteins; living cells were imaged 16 h p.t. (B) Images such as those in (A) were used to calculate the % nucleoli with punctate localization of wt M protein (analysis of ≥ 99 nucleoli for each condition; results are representative of 3 independent experiments).

To determine whether HeV M interaction with Treacle-enriched puncta results in gross effects on their structure or distribution, we employed super-resolution microscopy using single molecule localizations (*d*STORM)^30,31^ to analyze cells immunostained for Treacle. This achieved spatial resolution at least as good as 40 nm in *xy* and 80 nm in *z*, enabling, to our knowledge, the first super-resolution measurement of the dimensions of Treacle-enriched puncta. 3D super-resolution images indicated that Treacle-enriched puncta are largely spheroidal structures (Fig. 6A), with axial cross-sections similar to the feature size detected in 2D super-resolution images (Fig. 6B), for which the mean area of puncta was 0.077 µm^2^ (n = 1959 puncta in 226 nucleoli, 59 cells). Importantly, the mean area was not significantly affected by expression of wt or mutated GFP-HeV M (Fig. 6C), compared with expression of GFP alone. Furthermore, there was no difference in the mean number of nucleoli per cell (Fig. 6D), or puncta per nucleolus (Fig. 6E) detected by *d*STORM. Thus, HeV M protein does not appear to affect either the number of Treacle-enriched puncta or their dimensions, such that any functional effect of HeV M protein punctate localization is likely to result from specific intra-nucleolar protein interactions.

**Figure 6.**
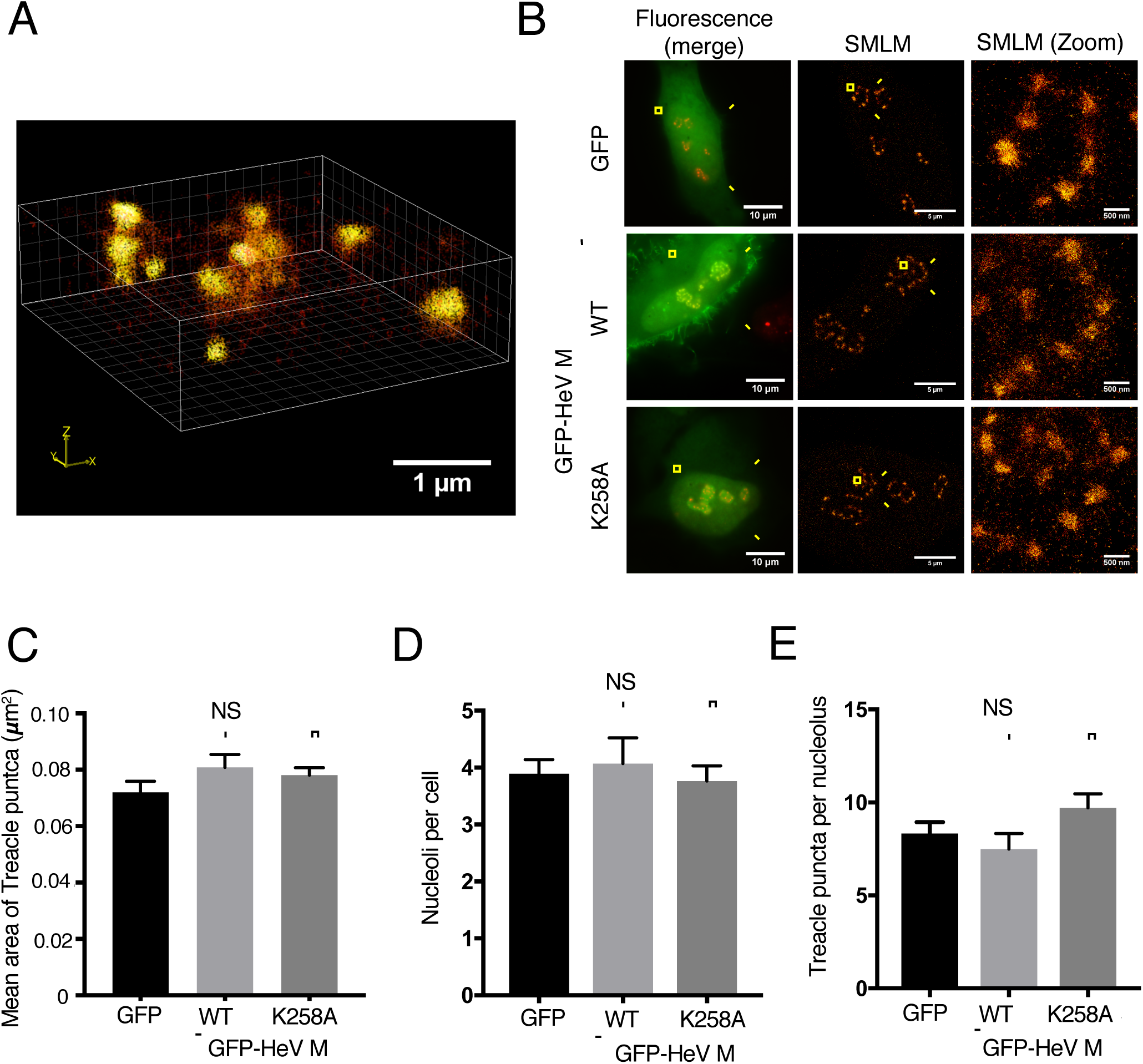
Localization of HeV M to puncta does not impact gross structure or number of puncta. (A,B) HeLa cells expressing GFP alone (A) or the indicated proteins (B) were fixed and immunolabeled for Treacle before analysis: (A) *d*STORM single molecule localization microscopy (SMLM) was used to generate 3D images of Treacle punct; (B) Fluorescence microscopy was used to detect GFP and Treacle (red/yellow in merged image), followed by SMLM to detect Treacle in GFP positive cells. (C-E) Images such as those shown in Fig. 6B were used to determine (C) the number of nucleoli per cell (mean ± SEM, n ≥ 14 cells), (D) the number of Treacle puncta per nucleolus (mean ± SEM, n ≥ 57 nucleoli), and (E) the area of Treacle puncta (mean ± SEM, n ≥ 427 puncta). Statistical analysis used Student’s t-test; NS = not significant.

### HeV M protein inhibits rRNA biogenesis by hijacking the DDR-Treacle pathway

Treacle regulates rRNA production and is required for efficient ‘basal’ synthesis of rRNA, providing a molecular target for cellular regulation of rRNA, as suggested by its role in the DDR^2,4,7^. To examine whether M protein targets puncta to exploit such a regulatory mechanism, we assessed the effect of HeV infection on rRNA synthesis, using *in situ* detection of rRNA in the nucleolus by the Click-iT^TM^ RNA Imaging Kit, which was previously used to identify the role of Treacle in the DDR^4^. A significant (p < 0.0001) reduction in rRNA synthesis was observed in HeV-infected cells compared with mock-infected cells (Fig. 7A, B) indicating that HeV can inhibit Treacle function. Expression of GFP-HeV M alone also significantly (p < 0.001) reduced rRNA synthesis, with the extent of reduction (c. 35%, Fig. 7C, D) similar to that reported following knockdown of Treacle^4^. Notably, treatment of cells with actinomycin D (ActD), which arrests rRNA synthesis globally by inhibiting RNA Polymerase I (RNA Pol I), reduced rRNA production to a greater extent (> 90% reduction of rRNA synthesis, Fig. 7D). Thus, the effect of HeV infection and HeV M protein expression is consistent with specific inhibition of Treacle-dependent processes. Consistent with this idea, no effect on rRNA production was observed following expression of GFP-HeV M-K258A, which does not bind to Treacle (Fig. 7C, D).

**Figure 7.**
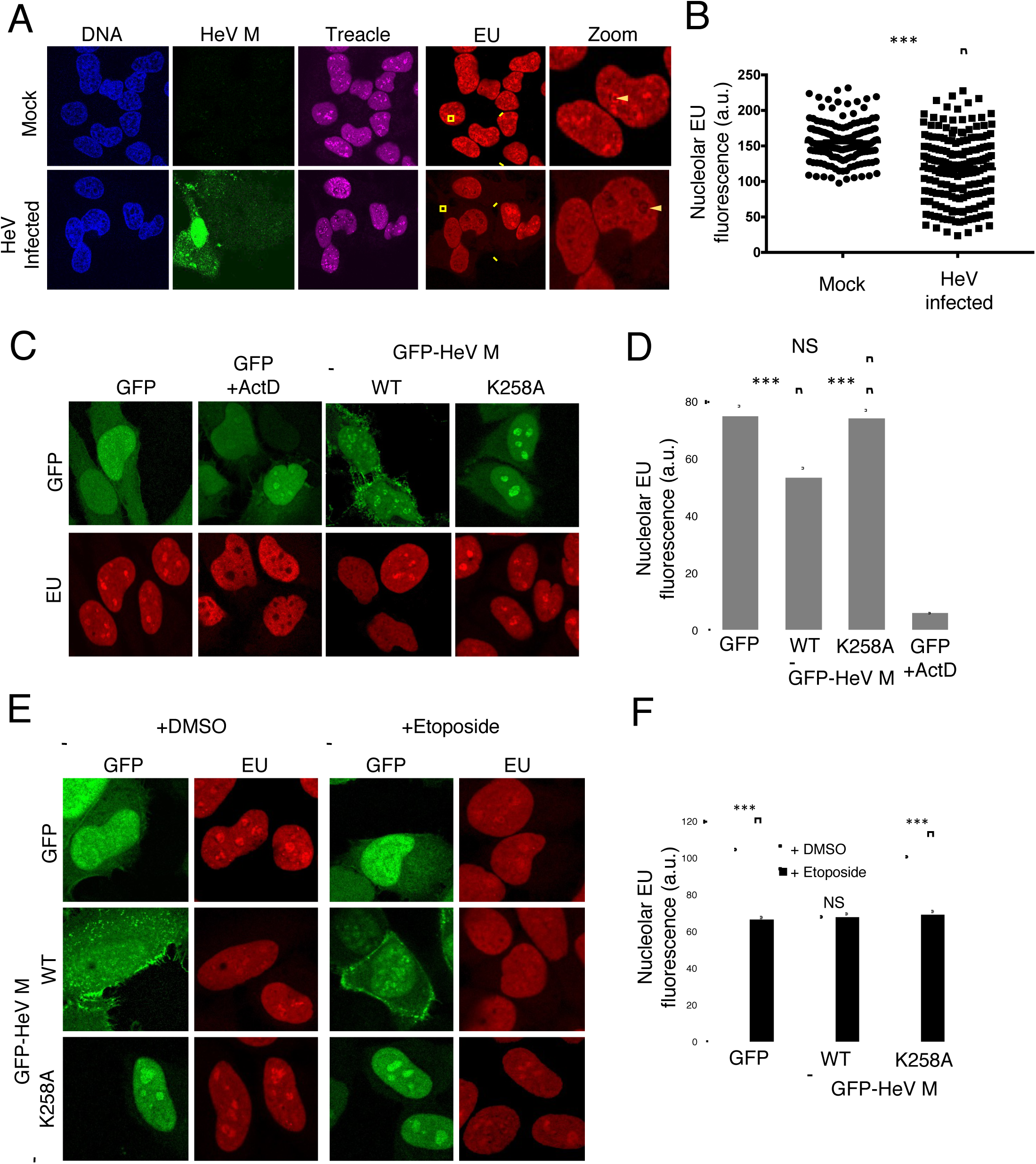
HeV and HeV M protein inhibit rRNA production. rRNA production in HeLa cells was assayed by labelling nascent RNA using EU/Alexa 594 for analysis by CLSM; cell samples analyzed were: (A, B) infected with HeV, fixed at 24 h p.i., and labelled with anti-HeV M protein and anti-Treacle antibodies, and Hoechst 33342 (DNA) (≥ 156 nucleoli analyzed for each condition over 2 biological replicates; arrows indicate nucleoli); (C, D) transfected to express the indicated GFP-fused proteins before treatment with or without ActD and fixation at 16 h p.t. (histogram shows mean nucleolar EU fluorescence ± SEM, n ≥ 31 nucleoli; data from a single assay, representative of 6 independent assays); (E, F) transfected to express the indicated proteins before treatment without (DMSO) or with etoposide for 3 h, and fixation at 16 h p.t. (mean nucleolar EU fluorescence ± SEM, n ≥ 39 nucleoli; data from a single assay, representative of 3 independent assays).

Since the effect of HeV M protein on rRNA biogenesis was consistent with that observed during a DDR^4^, we sought to define whether the same molecular pathway is being utilized by testing the impact of HeV M protein on the inhibition of rRNA synthesis induced by DNA damage. Using etoposide, we confirmed that DNA damage inhibits rRNA biogenesis, and found that it does so to a similar extent as HeV M protein expression (Fig. 7E, F). Importantly, combination of these stimuli produced no additional effect, suggesting that HeV M exploits the same pathway of rRNA silencing that occurs during the DDR, which is Treacle-dependent^2,4^ (Fig. 7E, F). To confirm directly the role of Treacle in HeV M protein-mediated suppression of rRNA biogenesis, we analyzed rRNA synthesis in cells transfected with siRNA to knockdown Treacle expression and/or transfected to express HeV M protein, detecting no additive effect by combining these stimuli (Fig. 8A), confirming the effect is Treacle-dependent.

**Figure 8.**
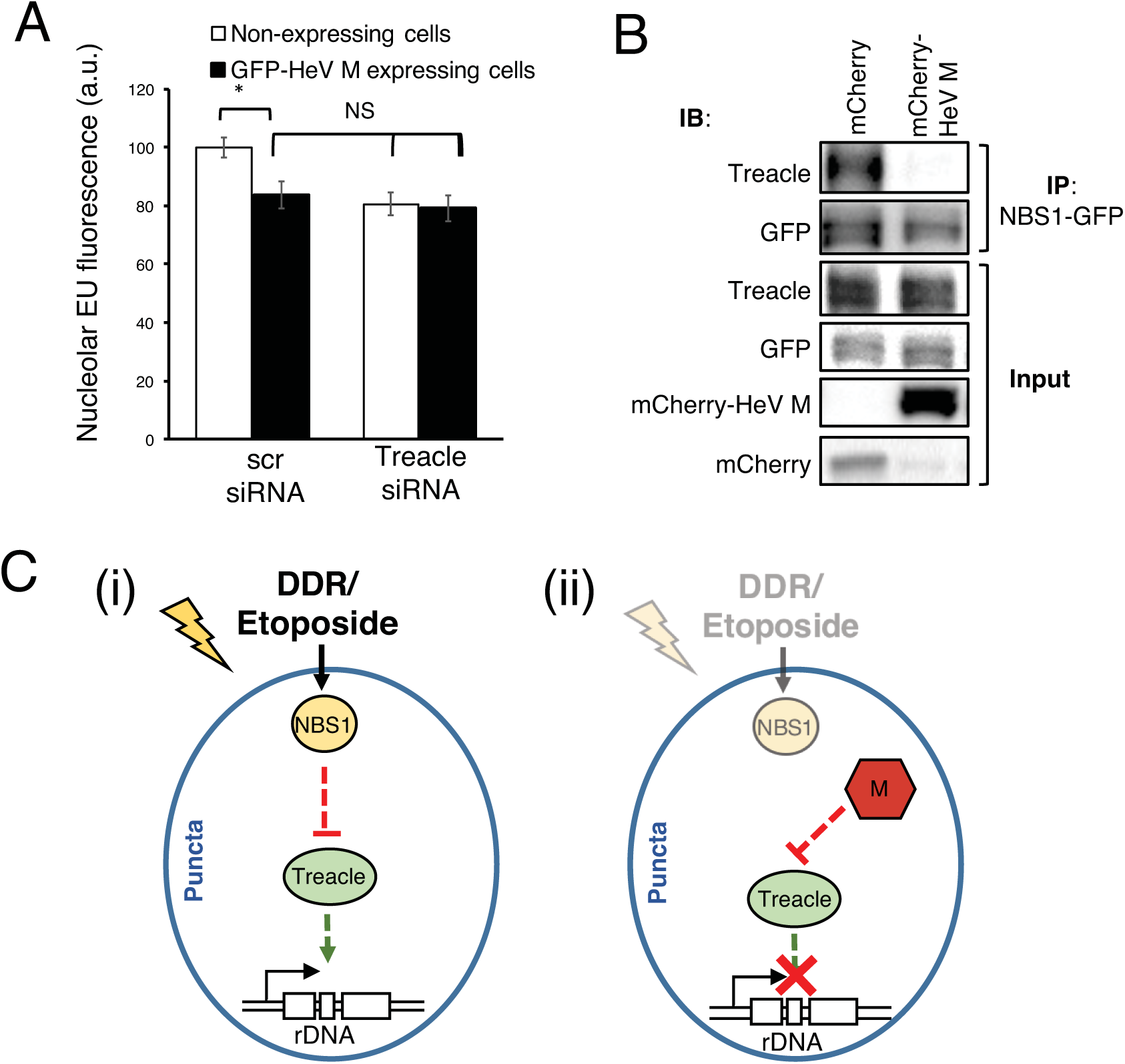
HeV M protein inhibits rRNA synthesis by a Treacle-dependent mechanism and disrupts the Treacle-NBS1 complex. (A) HeLa cells were transfected with scrambled (scr) or Treacle siRNA followed by transfection to express GFP-HeV M protein before fixation at 16 h p.t. and processing to label nascent rRNA for analysis by CLSM as described in the legend to Fig. 7 (histogram shows mean nucleolar EU fluorescence ± SEM, n ≥ 23 nucleoli; data from a single assay representative of 3 independent assays). (B) HEK-293T cells co-transfected to express the indicated mCherry protein with NBS1-GFP were subjected to IP for NBS1-GFP followed by analysis by IB using the indicated antibodies; results are representative of 3 independent experiments. (C) Models for DDR (i) and HeV M protein (ii)-mediated inhibition of rRNA synthesis. Treacle is required for basal rRNA synthesis and localizes to Treacle-enriched subnucleolar puncta. During the DDR (i), Treacle function is inhibited by NBS1 with which Treacle forms a complex. During infection (ii), HeV M localizes to Treacle-enriched puncta and binds to Treacle resulting in inhibition of Treacle-dependent function independently of a DDR. M protein appears to compete with NBS1 for binding to Treacle (B), suggesting that it binds at or close to the Treacle-NBS1 interaction site to mimic a DDR activated NBS1-Treacle complex.

### HeV M protein disrupts Treacle-NBS1 complexes

The properties of HeV M protein with respect to Treacle are analogous to those of DDR-activated NBS1, including the dependence of punctate localization and rRNA-inhibitory function on expression of Treacle^2,4,7^. Since HeV M protein appears to appropriate the DDR-Treacle pathway, we reasoned that M protein might bind at the same or overlapping site(s) in Treacle as NBS1, thereby mimicking the DDR-activated response but in the absence of DNA-damage. Such competitive binding would be consistent with the lack of additional effects of DNA-damage on rRNA biosynthesis in M-protein expressing cells (above). To examine this we used IP to analyze Treacle-NBS1 complexes in cells expressing mCherry-HeV M or mCherry alone. Consistent with previous reports^4,32^, NBS1-Treacle complexes were readily detected by co-IP from control cells (Fig. 8B). However, the association was not detectable in IPs from HeV M protein-expressing cells, indicating that HeV M protein efficiently displaces NBS1 from the complex (Fig. 8B).

### RABV P3 protein interacts with Treacle and inhibits rRNA synthesis

The above data provide, to our knowledge, the first evidence of a specific intra-nucleolar function for a protein of a mononegavirus. To examine whether this mechanism might be utilized by nucleolar proteins of viruses of other families in this order, we used the P3 protein of the rhabdovirus RABV, which enters nucleoli and interacts with NCL^16^. IP analysis of cells expressing GFP-fused RABV P3 clearly indicated interaction of P3 with endogenous Treacle (Fig. 9A) and analysis of cells expressing P3 or a control protein (HeV M K258A) using the Click-iT^TM^ RNA Imaging Kit indicated that P3 significantly inhibits rRNA biogenesis (Fig. 9B).

**Figure 9.**
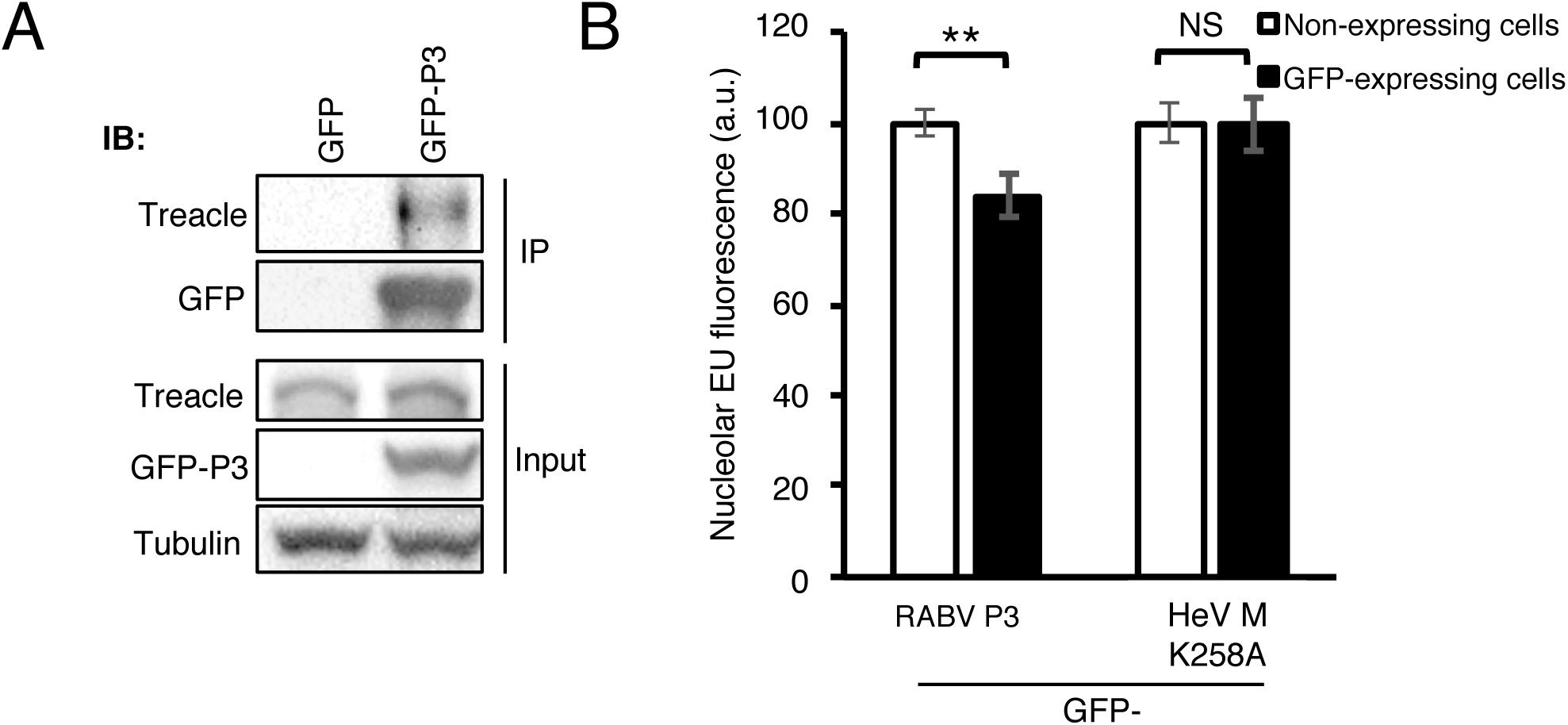
RABV P3 protein interacts with Treacle and inhibits rRNA synthesis. (A) HEK-293T cells expressing the indicated GFP proteins were subjected to IP for GFP before analysis by IB using the indicated antibodies; results are representative of 2 independent experiments. (B) HeLa cells were transfected to express GFP-RABV-P3 or GFP-HeV M-K258A before fixation at 16 h p.t. and processing to label nascent rRNA for analysis by CLSM as described in the legend to Fig. 7 (histogram shows mean nucleolar EU fluorescence intensity ± SEM, n ≥ 28 nucleoli; data from a single assay representative of 2 independent assays).

## DISCUSSION

In this study, we identified the rRNA transcriptional regulator Treacle as a novel target for subversion of the host cell by a microbial pathogen. Treacle is a critical component in cellular mechanisms to regulate rDNA transcription following DNA damage^2,4^. The role of Treacle in the DDR appears to be due to interaction with NBS1, an integral player in the DDR^2,4^. Current data on the NBS1-Treacle complex are consistent with a model whereby DDR-activated NBS1 inhibits rRNA biogenesis by suppressing Treacle’s function in maintaining basal levels of rRNA (Fig. 8C(i)). Following the finding that viral proteins can bind to Treacle (Fig. 4), we hypothesized that such a mechanism could be exploited by viruses through the formation of a complex with Treacle analogous to that formed by NBS1, thereby inducing the DDR pathway without the requirement for a DNA damage signal (Fig. 8C(ii)). Consistent with this, our data indicate that M protein interaction with Treacle displaces NBS1 from the complex, and appropriates the downstream pathway that would be activated during a DDR. These data indicate that binding of specific proteins to Treacle can directly inhibit rRNA biogenesis, consistent with the idea that specific regulation of Treacle function (Fig. 8C(i)), rather than some alternative function of nucleolar NBS1 or of the NBS1-Treacle complex, is responsible for rRNA suppression in the DDR. Notably, one prediction of our model (Fig. 8C (ii)) is that NBS1 would have antiviral properties and, consistent with this, a previous genome-wide functional genomics screen of HeV infection identified NBS1 as antiviral (Z-score = 2.77)^15^.Thus, our data support both the model for DDR mediated rRNA silencing *via* direct inhibition of Treacle-dependent rRNA biogenesis, and the proposed novel mechanism for viral subversion of this process.

The precise mechanisms underlying Treacle function in rRNA production are not fully resolved, but likely involve interaction with other nucleolar components critical to rDNA transcription such as RNA Pol I and the RNA Pol I transcription factor UBF1, with which Treacle co-localizes, as well as direct interaction of Treacle with rDNA^7,33^. Intriguingly, our IP/MS analysis suggested that HeV M protein interactors within the puncta include UBF1, indicating that HeV M protein might form complexes incorporating UBF1 and Treacle. Consistent with this, UBF1 was previously identified as an interactor of NiV M protein, although the subnucleolar localization of the interaction was not elucidated^22^. The interactions of Treacle, RNA Pol I, UBF1 and rDNA appear to occur within Treacle-enriched puncta, suggesting that this subnucleolar localization facilitates specific interaction network(s) enabling roles in rRNA production and stress responses^2,4^, which our data now indicate are targeted by viruses. Delineation of the specific signaling pathways/networks that underpin the diverse functions of nucleoli and nucleolar compartments is hampered by their molecular complexity. Based on the findings of the current study with respect to NBS1-Treacle interaction, it is likely that further analysis of virus-nucleolar targeting will provide additional insights into the organization and functions of Treacle-enriched puncta, particularly given the availability of mutated viral proteins that differ in their specific interactions with these structures.

Other than roles in rDNA transcription, Treacle also functions in pre-rRNA processing suggesting that the DDR pathway induced by DNA damage or by viral proteins is likely to have important effects on rRNA and ribosome function beyond suppression of rRNA synthesis. Notably, the roles of Treacle in pre-rRNA processing involve association with the Box C/D small nucleolar ribonucleoprotein (snoRNP) complex. The Box C/D snoRNP complex methylates pre-rRNA and is composed of four core proteins: FBL, NOP56, NOP58 and NHP2L1^34^. Three of these proteins (FBL, NOP56 and NOP58) have been implicated as being important to a henipavirus infection in an siRNA screen, and confirmed or proposed to associate with M protein, although no direct function for these interactions has yet been demonstrated^15,22,23^. Our analysis indicates that FBL and HeV M protein do not colocalize within the Treacle-enriched puncta (Fig. 2), and FBL interaction is unaffected by K258A (Fig. S3). Thus, the interaction of M protein with FBL appears distinct from the role of HeV M protein in Treacle-dependent modulation of rRNA production. Taken together with the findings that other nucleolar interactors including NPM1 and NCL were unaffected by K258A mutation (Fig. 3C), that IP/MS analysis of henipavirus M protein has identified multiple additional nucleolar interactors (our data and others^22,23^), and that a functional genomics screen identified several nucleolar factors able to inhibit infection^15^, this suggests that M protein is likely to have additional nucleolar roles additional to the Treacle/puncta interaction. Finally, our observation that RABV P3, an isoform of the P protein that is distinct from the M protein, targets Treacle (in addition to NCLref.) and inhibits rRNA synthesis suggests that the same functions have developed independently in viruses of different families, indicative of broad importance^16^. Taken together, these data indicate that the nucleolus represents an important and potentially diverse interface for mononegaviruses.

## Methods

### Antibodies and Reagents

Rabbit anti-TCOF1/treacle was purchased from Proteintech (Cat #11003-1-AP) and used for immunofluorescence (IF) and immunoblotting (IB). Mouse anti-GFP (Sigma-Aldrich, Cat #11814460001 ROCHE) was used for IB. Mouse anti-HeV M (Generated at AAHL^35^ was used for IF, as previously described^21^. Anti-FBL (Abcam, Cat #Ab4566) was used for IF and immunoblotting. Secondary antibodies used for IF (goat anti-rabbit Alexa Fluor 568 (Cat #A-11011), goat anti-rabbit Alex Fluor 647 (Cat #A-21245)) were purchased from Thermo Fisher Scientific. Secondary antibodies used for IB (goat anti-rabbit (Cat #AP307P) and goat anti-mouse (Cat #AP308P) IgG horse radish peroxidase (HRP)-conjugated-antibodies) were purchased from Merck. rRNA synthesis was inhibited by treating cells with Actinomycin D (ActD) (Thermo Fisher Scientific, Cat #11805017) for 1 h at 500 ng/mL. DNA damage was induced by treating cells with 50 µM etoposide for 4 h.

### Cell Culture

HeLa and HEK-293T cells were maintained in Dulbecco’s Modified Eagle Medium (DMEM, ThermoFisher Scientific, Cat# 11965092) supplemented with 10% Fetal Calf Serum (FCS) at 37°C, 5% CO_2_.

### Transfections

Plasmids for expression in mammalian cells of HeV-M protein (Accession Number AEB21196.1), NiV-M (Accession Number AAY43914.1), and mutants thereof, fused at the N terminus to GFP, were generated by directional cloning of M gene cDNA into the multiple cloning site of the pEGFP-C1 vector, as previously described^27^. The RV CVS strain protein, P3, was fused at the N-terminus to GFP (GFP-RV-P3) and has been described previously^16^. pmRFP-N1-Fibrillarin, mCherry-NCL and NBS1-GFP were kind gifts from P. Roussel (Université Paris Diderot), Keiichi I. Nakayama (Kyushu University) and S. Elledge (Harvard University), respectively. siRNA targeting Treacle consisted of a pool of 3 treacle-specific siRNAs, synthesized by Bioneer Pacific^®^ (Sequences (5’-3’): GGUCUCCAUCCAAGGUGAAA(dTdT); CAGUAGUGAGGAGUCAUCA(dTdT); GCAAGCUAAGAAAACCCGU(dTdT)).

Plasmids were transfected into HEK-293T cells and HeLa cells using Lipofectamine 2000^TM^ and Lipofectamine 3000^TM^, respectively, according to the manufacturer’s instructions (Thermo-Fisher Scientific). siRNA (100 nM final) was transfected into HeLa cells using DharmaFECT 1 Transfection Reagent^TM^ (GE Dharmacon) according to the manufacturer’s instructions.

### Confocal Laser Scanning Microscopy and image analysis

Standard CLSM used a Leica SP5 or Nikon C1 microscope with 60x oil immersion objective (NA 1.4), and a heated chamber (37°C) for live-cell analysis. Image analysis was performed using ImageJ freeware software. To generate super-resolution images of living cells we used a Leica SP8 with Hyvolution. Hyvolution works by narrowing the pinhole to increase the effective resolution of the system (0.5 airy units was used in these experiments). Images were acquired with 4 x line averages using high sensitivity HyD detectors and then deconvolved using Huygens (SVI, Netherlands). For IF staining, cells seeded onto glass coverslips were fixed with 4% paraformaldehyde (37°C, 10 min), permeabilized with 0.25% Triton X-100 (room temperature (RT), 5 min), and blocked with 1% bovine serum albumin (BSA) in PBS (RT, 1 h), before primary and secondary antibody labeling (RT, 90 min each), and coverslips were mounted onto glass slides with Mowiol.

### *d*STORM imaging and analysis

For *d*STORM analysis, HeLa cells were fixed 24 h p.t. using 4% paraformaldehyde (37ºC, 10 min), and permeabilized using 0.1% Triton X-100 (RT, 10 min). Slides were then blocked with 2% BSA in PBS (RT, 30 min) before labelling using anti-Treacle (3 µg/ml, 1 h, RT) and Alexa Fluor-647-conjugated anti-rabbit secondary antibodies (5 µg/ml, 45 min, RT). Labelled cells were imaged in a switching buffer of 10% glucose, 100 mM mercaptoethylamine, 400 µg/ml glucose oxidase and 35 µg/ml bovine catalase in PBS, made to pH 8.5 using 1M KOH. Imaging used a home-built super-resolution set-up based on a previously described system^31^ (details in Supplemental Material and Methods). Images were generated using rapidSTORM, and analysis to determine puncta area used ImageJ after first smoothing 2D super-resolution images with a Gaussian Blur (r = 0.5) to account for localization precision error (see Supplemental Materials for details).

For 3D imaging, a cylindrical lens (f = 1000 mm) was inserted into the imaging optical path and calibrated by scanning (in z) a 0.1 µm Tetraspeck sphere embedded in a water soluble gel matrix and fitting the changing lateral distortions of emission intensity with a 2D Gaussian as a function of axial position^36^. 3D *d*STORM of cells was achieved with the calibration file in rapidSTORM to localize 3D coordinates of single molecules into a text file that was used to plot 3D models in ViSP^37^. Puncta were color-coded for relative 3D density to highlight spheroidal conformation.

### Co-immunoprecipitation (co-IP)

All co-IPs were performed in 6 cm dishes of HEK-293T cells that were transfected to express the indicated proteins and lysed at 24 h p.t. with Lysis Buffer (10 mM Tris/Cl pH 7.5; 150 mM NaCl; 0.5 mM EDTA; 0.5% NP-40, 1x Protease Inhibitor Cocktail (PIC; Sigma-Aldrich Cat #11697498001)) for 30 min at 4°C. Supernatants were collected by centrifugation at 20,000x g for 15 min at 4°C and 10% of the cleared lysate was collected for ‘input’ analysis; the remaining lysate was subjected to IP using 10 µL of GFP-Trap^®^ beads (Chromotek) as previously described^38^. Beads were washed 3 times with Dilution Buffer (10 mM Tris/Cl pH 7.5; 150 mM NaCl; 0.5 mM EDTA, 1x PIC). Samples intended for IB analysis were re-suspended in 2x SDS-PAGE sample loading buffer. Samples intended for mass-spectrometry analysis were subjected to a final wash containing no PIC.

### Mass Spectrometry and analysis

Processing of co-IP samples was performed at the Bio21 Mass Spectrometry and Proteomics facility (University of Melbourne). Proteins bound to beads from IP assays were eluted using trifluoroacetic acid (TFA) prior to readjustment of pH to ∼8 with triethylammonium bicarbonate buffer (TEAB) and trypsin digestion. LC MS/MS was performed on a QExactive plus Orbitrap mass spectrometer (Thermo Scientific) with a nanoESI interface in conjunction with an Ultimate 3000 RSLC nanoHPLC (Dionex Ultimate 3000). The LC system was equipped with an Acclaim Pepmap nano-trap column (Dinoex-C18, 100 Å, 75 µm x 2 cm) and an Acclaim Pepmap RSLC analytical column (Dinoex-C18, 100 Å, 75 µm x 50 cm). Result files were searched against the SwissProt database in a target decoy fashion using MASCOT (Version 2.4.1, Matrix Science, UK). Details of the mass-spectrometry procedure and analysis are provided in Supplemental Materials and Methods. Interactions identified were confirmed in ≥ 2 separate assays.

### SDS-PAGE and Immunoblotting

IP elutions and input samples from co-IP assays were separated on a 10% denaturing gel by SDS-PAGE before transfer to a nitrocellulose membrane using a BioRad Trans-Blot semi-dry apparatus. After blocking (5% non-fat milk in PBS with 1% Tween20 (PBST)), the membranes were incubated with primary antibodies followed by HRP-conjugated goat anti-rabbit or anti-mouse secondary antibodies, and imaged on a Gel Doc™ XR+ Gel Documentation System.

### 5-ethnyl uridine (EU) incorporation assays

Analysis of rRNA was performed as previously^4^. Briefly, detection of nascent rRNA was determined using the Click-iT^TM^ RNA Alexa Fluor 594 Imaging Kit (Thermo-Fisher, Cat# C10330). Cells were incubated for 1 h in the presence of EU before fixation in 4% paraformaldehyde at RT for 12 min, and permeabilization in 0.25% Triton X-100 for 5 min at RT. Samples were then processed according to the manufacturer’s recommendations to label incorporated EU with Alexa Fluor 594. Cells were imaged by CLSM to detect labelling of nascent rRNA by measuring the fluorescence intensity of Alexa Fluor 594 within nucleoli. Quantitative analysis to determine EU fluorescence (arbitrary units, a.u.) was performed using ImageJ software, with nucleoli identified using differential interference contrast (DIC) microscopy and IF labeling of nucleoli by anti-FBL or anti-Treacle/TCOF1.

### Virus infections

All work with infectious virus was conducted at the CSIRO Australian Animal Health Laboratory (AAHL) in Biosafety Level (BSL)-4 laboratories. For analysis by IF microscopy, HeLa cells were seeded onto coverslips and mock- or HeV-infected (MOI 5) prior to fixation using 4% paraformaldehyde and permeabilization with 0.25% Triton X-100. IF labeling was performed using antibody to HeV M alone or together with antibody to Treacle, followed by Alexa Fluor^®^ 488 (for HeV M) and Alex Fluor^®^ 568 (for Treacle) secondary antibody. For EU analysis, HeLa cells were seeded into 8-well chambers and mock or HeV-infected (MOI 5). EU was added to media 1 h prior to fixation at 24 h p.i. with 4% paraformaldehyde, and analysis as described above. In all cases, cells were decontaminated for 2 h using 4% paraformaldehyde before removal from BSL-4 laboratories.

### Tissue Culture Infective Dose (TCID_50_) analysis

HeLa cells were transfected without siRNA, or with scr siRNA, Treacle siRNA or PLK1 siRNA (3 days) prior to infection with HeV at MOI 0.5. siRNA to PLK1, which induces apoptosis and cell death, was used as a transfection and indirect positive control as previously described^15^. Viral TCID_50_ was determined as described previously^39^. Briefly, samples were titrated in triplicate in 96-well plates and co-cultured with Vero cells for 3 days. The infectious titer was calculated by the method of Reed and Muench^40^.

## Acknowledgments

We acknowledge Cassandra David for assistance with tissue culture, and the facilities and technical assistance of Monash Micro Imaging (Monash University) and the Biological Optical Microscopy Platform (University of Melbourne). Mass spectrometry was performed at the Mass Spectrometry and Proteomics facility at the Bio21 Institute (University of Melbourne) and at the Monash Biomedical Proteomics Facility (Monash University). We thank Angus Lamond (University of Dundee) for helpful discussions and acknowledge his contributions to securing funding (Australian Research Council (ARC) grant DP150102569). This work was supported by: National Health and Medical Research Council Australia grants 1125704 & 1079211 to GWM, and 1044228 to HJN; ARC grant DP150102569 to GWM, LFW and Angus I. Lamond (University of Dundee); NRF-CRP grant NRF2012NRF-CRP001-056 to LFW.

## References

1 Ciccia, A. & Elledge, S. J. The DNA damage response: making it safe to play with knives. Mol Cell 40, 179–204, doi:10.1016/j.molcel.2010.09.019 (2010).

2 Ciccia, A. et al. Treacher Collins syndrome TCOF1 protein cooperates with NBS1 in the DNA damage response. Proc Natl Acad Sci U S A 111, 18631–18636, doi:10.1073/pnas.1422488112 (2014).

3 Kruhlak, M. et al. The ATM repair pathway inhibits RNA polymerase I transcription in response to chromosome breaks. Nature 447, 730–734, doi:10.1038/nature05842 (2007).

4 Larsen, D. H. et al. The NBS1-Treacle complex controls ribosomal RNA transcription in response to DNA damage. Nat Cell Biol 16, 792–803, doi:10.1038/ncb3007 (2014).

5 Larsen, D. H. & Stucki, M. Nucleolar responses to DNA double-strand breaks. Nucleic Acids Res 44, 538–544, doi:10.1093/nar/gkv1312 (2016).

6 Boisvert, F. M., van Koningsbruggen, S., Navascues, J. & Lamond, A. I. The multifunctional nucleolus. Nat Rev Mol Cell Biol 8, 574–585, doi:10.1038/nrm2184 (2007).

7 Valdez, B. C., Henning, D., So, R. B., Dixon, J. & Dixon, M. J. The Treacher Collins syndrome (TCOF1) gene product is involved in ribosomal DNA gene transcription by interacting with upstream binding factor. Proc Natl Acad Sci U S A 101, 10709–10714, doi:10.1073/pnas.0402492101 (2004).

8 Werner, A. et al. Cell-fate determination by ubiquitin-dependent regulation of translation. Nature 525, 523–527, doi:10.1038/nature14978 (2015).

9 Dixon, J., Trainor, P. & Dixon, M. J. Treacher Collins syndrome. Orthod Craniofac Res 10, 88–95, doi:10.1111/j.1601-6343.2007.00388.x (2007).

10 Sakai, D. & Trainor, P. A. Face off against ROS: Tcof1/Treacle safeguards neuroepithelial cells and progenitor neural crest cells from oxidative stress during craniofacial development. Dev Growth Differ 58, 577–585, doi:10.1111/dgd.12305 (2016).

11 Parlato, R. & Liss, B. How Parkinson’s disease meets nucleolar stress. Biochim Biophys Acta 1842, 791–797, doi:10.1016/j.bbadis.2013.12.014 (2014).

12 Derenzini, M., Montanaro, L. & Trere, D. Ribosome biogenesis and cancer. Acta Histochem 119, 190–197, doi:10.1016/j.acthis.2017.01.009 (2017).

13 Rawlinson, S. M. & Moseley, G. W. The nucleolar interface of RNA viruses. Cell Microbiol 17, 1108–1120, doi:10.1111/cmi.12465 (2015).

14 Salvetti, A. & Greco, A. Viruses and the nucleolus: the fatal attraction. Biochim Biophys Acta 1842, 840–847, doi:10.1016/j.bbadis.2013.12.010 (2014).

15 Deffrasnes, C. et al. Genome-wide siRNA Screening at Biosafety Level 4 Reveals a Crucial Role for Fibrillarin in Henipavirus Infection. PLoS Pathog 12, e1005478, doi:10.1371/journal.ppat.1005478 (2016).

16 Oksayan, S. et al. Identification of a role for nucleolin in rabies virus infection. J Virol 89, 1939–1943, doi:10.1128/JVI.03320-14 (2015).

17 Wang, Y. E. et al. Ubiquitin-regulated nuclear-cytoplasmic trafficking of the Nipah virus matrix protein is important for viral budding. PLoS Pathog 6, e1001186, doi:10.1371/journal.ppat.1001186 (2010).

18 Wang, L. F., Mackenzie, J. S. & Broder, C. C. in Fields Virology Vol. 2 (ed D.M.; Howley Knipe, P.M.) 286–313 (Lippincott Williams & Wilkins, 2013).

19 Liljeroos, L. & Butcher, S. J. Matrix proteins as centralized organizers of negative-sense RNA virions. Front Biosci (Landmark Ed) 18, 696–715 (2013).

20 Watkinson, R. E. & Lee, B. Nipah virus matrix protein: expert hacker of cellular machines. FEBS Lett 590, 2494–2511, doi:10.1002/1873-3468.12272 (2016).

21 Monaghan, P. et al. Detailed morphological characterisation of Hendra virus infection of different cell types using super-resolution and conventional imaging. Virol J 11, 200, doi:10.1186/s12985-014-0200-5 (2014).

22 Pentecost, M. et al. Evidence for ubiquitin-regulated nuclear and subnuclear trafficking among Paramyxovirinae matrix proteins. PLoS Pathog 11, e1004739, doi:10.1371/journal.ppat.1004739 (2015).

23 Sun, W. et al. Matrix proteins of Nipah and Hendra viruses interact with beta subunits of AP-3 complexes. J Virol 88, 13099–13110, doi:10.1128/JVI.02103-14 (2014).

24 Audsley, M. D., Jans, D. A. & Moseley, G. W. Roles of nuclear trafficking in infection by cytoplasmic negative-strand RNA viruses: paramyxoviruses and beyond. J Gen Virol 97, 2463–2481, doi:10.1099/jgv.0.000575 (2016).

25 Hutten, S. et al. An intranucleolar body associated with rDNA. Chromosoma 120, 481–499, doi:10.1007/s00412-011-0327-8 (2011).

26 Lam, Y. W. & Trinkle-Mulcahy, L. New insights into nucleolar structure and function. F1000Prime Rep 7, 48, doi:10.12703/P7-48 (2015).

27 McLinton, E. C. et al. Nuclear localization and secretion competence is conserved amongst henipavirus matrix proteins. J Gen Virol, doi:10.1099/jgv.0.000703 (2017).

28 Emmott, E. & Hiscox, J. A. Nucleolar targeting: the hub of the matter. EMBO Rep 10, 231–238, doi:10.1038/embor.2009.14 (2009).

29 Isaac, C. et al. Characterization of the nucleolar gene product, treacle, in Treacher Collins syndrome. Mol Biol Cell 11, 3061–3071 (2000).

30 Brice, A. et al. Quantitative Analysis of the Microtubule Interaction of Rabies Virus P3 Protein: Roles in Immune Evasion and Pathogenesis. Sci Rep 6, 33493, doi:10.1038/srep33493 (2016).

31 Whelan, D. R., Holm, T., Sauer, M. & Bell, T. D. M. Focus on Super-Resolution Imaging with Direct Stochastic Optical Reconstruction Microscopy (dSTORM). Aust J Chem 67, 179–183, doi:10.1071/Ch13499 (2014).

32 Sakai, D., Dixon, J., Achilleos, A., Dixon, M. & Trainor, P. A. Prevention of Treacher Collins syndrome craniofacial anomalies in mouse models via maternal antioxidant supplementation. Nat Commun 7, 10328, doi:10.1038/ncomms10328 (2016).

33 Lin, C. I. & Yeh, N. H. Treacle recruits RNA polymerase I complex to the nucleolus that is independent of UBF. Biochem Biophys Res Commun 386, 396–401, doi:10.1016/j.bbrc.2009.06.050 (2009).

34 Watkins, N. J. & Bohnsack, M. T. The box C/D and H/ACA snoRNPs: key players in the modification, processing and the dynamic folding of ribosomal RNA. Wiley Interdiscip Rev RNA 3, 397–414, doi:10.1002/wrna.117 (2012).

35 White, J. R. et al. Location of, immunogenicity of and relationships between neutralization epitopes on the attachment protein (G) of Hendra virus. J Gen Virol 86, 2839–2848, doi:10.1099/vir.0.81218-0 (2005).

36 Proppert, S. et al. Cubic B-spline calibration for 3D super-resolution measurements using astigmatic imaging. Opt Express 22, 10304–10316, doi:10.1364/OE.22.010304 (2014).

37 El Beheiry, M. & Dahan, M. ViSP: representing single-particle localizations in three dimensions. Nat Methods 10, 689–690, doi:10.1038/nmeth.2566 (2013).

38 Wiltzer, L. et al. Conservation of a unique mechanism of immune evasion across the Lyssavirus genus. J Virol 86, 10194–10199, doi:10.1128/JVI.01249-12 (2012).

39 Stewart, C. R. et al. Promotion of Hendra virus replication by microRNA 146a. J Virol 87, 3782–3791, doi:10.1128/JVI.01342-12 (2013).

40 Reed, L. & Muench, H. A simple method of estimating fifty percent endpoints. Am J Hygiene 27, 493–497 (1938).

41 Wolter, S. et al. rapidSTORM: accurate, fast open-source software for localization microscopy. Nat Methods 9, 1040–1041, doi:10.1038/nmeth.2224 (2012).

